# Differential contributions of Ca_V_2.2, GIRK, and HCN channel to the modulation of excitability by α-conotoxin Vc1.1 and baclofen in somatic and visceral sensory neurons

**DOI:** 10.1101/2025.03.21.644483

**Authors:** Mariana Brizuela, Anuja R. Bony, Sonia Garcia Caraballo, David J. Adams, Stuart M. Brierley

## Abstract

Chronic visceral pain is a key symptom of irritable bowel syndrome (IBS). Modulation of voltage-dependent calcium and potassium channels by G protein-coupled receptors (GPCRs) plays a key role in dampening nociceptive transmission. Baclofen and the analgesic peptide α-conotoxin Vc1.1 both activate GABA_B_ receptors (GABA_B_R), resulting in the inhibition of Ca_V_2.2 and Ca_V_2.3 calcium channels to reduce colonic nociception. Recent studies have also shown that GABA_B_R activation potentiates GIRK1/2 potassium channels in mammalian sensory afferent neurons. In this study, we investigated the expression of these ion channel targets in rodent and human dorsal root ganglion (DRG) neurons, including those innervating the colon. We also examined how Ca_V_2.2 and GIRK channel antagonists, as well as a GIRK channel activator, affect the passive and active electrical properties of adult mouse DRG neurons. Additionally, we assessed the effects of α-conotoxin Vc1.1 on neuronal excitability in the presence of the selective Ca_V_2.2 antagonist ω-conotoxin CVIE and the GIRK channel activator ML297. We further evaluated the impact of the GIRK channel antagonist Tertiapin-Q on excitability in mouse colonic DRGs and colonic afferents and explored the role of hyperpolarization-activated cyclic nucleotide-gated (HCN) channels in regulating membrane excitability of colonic DRGs. Our findings demonstrate that both Ca_V_2.2 inhibition and GIRK channel potentiation reduce excitability in mouse DRGs, likely mediating the analgesic effects of Vc1.1 and baclofen observed *in vivo*. However, our findings indicate that GIRK channel potentiation appears to play a limited role in modulating excitability in colon-innervating DRGs and colonic afferents. These findings suggest that neurons innervating different regions of the body employ distinct mechanisms to regulate neuronal excitability and nociceptive signaling.

**KEY POINTS SUMMARY:**

- GABA_B_R1, Ca_V_2.2, and GIRK1 are highly expressed in the thoracolumbar dorsal root ganglia (DRGs) of both mice and humans.
- In mouse DRGs, Ca_V_2.2 inhibition and GIRK channel potentiation contribute to reduced neuronal excitability.
- The analgesic peptide, α-conotoxin Vc1.1 reduces neuronal excitability by inhibiting Ca_V_2.2 and potentiating GIRK channels.
- However, potentiation of GIRK channels does not significantly affect the excitability in colon-innervating DRG neurons or colonic afferents.
- Sensory neurons innervating different body regions utilize distinct mechanisms to regulate their excitability.

## INTRODUCTION

Chronic pain is a major global health issue affecting more than 30% of the population (1). Chronic visceral pain is a particularly common form of chronic pain and is a key symptom of Irritable bowel syndrome (IBS), a chronic gastrointestinal disorder affecting 7-21% of individuals worldwide (2, 3). Although the pathophysiology of IBS is complex and not fully understood, substantial evidence indicates pain is driven by hypersensitivity of gut-innervating afferents and dysfunction along the gut-brain axis (4, 5). Peripheral hypersensitivity appears to be triggered by inflammatory or immune mediators, which contribute to the development of chronic visceral hypersensitivity (CVH) and neuroplastic changes within sensory pathways (4–6). These changes ultimately lead to heightened pain perception in the gastrointestinal tract (4, 5). Up-regulation of multiple ion channels and receptors in sensory afferents has been linked to the development of CVH in animal models (7–14). Consequently, targeting these ion channels and receptors may represent a promising therapeutic strategy for managing chronic visceral pain in IBS and chronic pain more broadly (5, 15–18).

α-Conotoxins are small, disulfide-rich peptides, typically 12−30 amino acids in length, derived from the venom of marine cone snails. These peptides are promising drug candidates for the treatment of chronic pain, and chronic visceral pain, due to their ability to selectively target a wide range of membrane receptors and ion channels (19–21). In particular, α-conotoxin Vc1.1 has demonstrated anti-nociceptive effects *in vitro* and anti-hyperalgesic actions in various *in vivo* models of neuropathic pain (20, 22, 23). Historically, the analgesic effects of Vc1.1 were attributed to its inhibitory action on neuronal nicotinic acetylcholine receptors (nAChRs) (24, 25). However, subsequent research has shown that its analgesic properties are primarily mediated through the activation of the G protein-coupled γ-aminobutyric acid type B receptor (GABA_B_R) (19, 24, 26, 27). GABA_B_Rs are broadly expressed on sensory neurons in both humans and rodents, including those innervating the gut (8). Our previous studies demonstrated that Vc1.1, along with its modified and cyclized form (cVc1.1), inhibits colonic sensory afferents, with these inhibitory effects being further amplified in an animal model of IBS (8, 28–30). Importantly, we identified that the mechanism of action of Vc1.1 involves the activation of GABA_B_Rs on colonic afferents, leading to downstream inhibition of voltage-gated calcium channels Ca_V_2.2 and Ca_V_2.3 and a reduction in the excitability of dorsal root ganglion (DRG) neurons innervating the mouse colon (8, 29). Furthermore, we have shown that native human DRG neurons and human pluripotent stem cell (hPSC)-derived sensory neurons are inhibited by the GABA_B_R agonists, baclofen and Vc1.1 (8, 31). Notably, the latter study confirmed the expression of G protein-coupled inwardly rectifying potassium (GIRK) channels in hPSC-derived sensory neurons. This is significant, as GABA_B_R activation is known to reduce neuronal excitability through inhibition of Ca_V_ channels, but also by potentiating GIRK channels (32).

In this study, we investigated the role of GIRK channels in modulating neuronal excitability in mouse DRG neurons, including those specifically innervating the colon. We assessed the contributions of Ca_V_2.2 and GIRK channels to the passive and active electrical properties of adult mouse DRG neurons. Additionally, we examined whether inhibitory effects of Vc1.1 on neuronal excitability are mediated by GABA_B_R activation, leading to downstream modulation of Ca_V_2.2 and GIRK1/2 channels. Finally, we explored the roles of GIRK channels and hyperpolarization-activated cyclic nucleotide-gated (HCN) channels in regulating the excitability of DRG neurons innervating the colon, as well as colonic afferents, to further elucidate the mechanisms by which α-conotoxins modulate colonic neuronal excitability.

## MATERIALS AND METHODS

### Animals

All animal procedures were conducted in accordance with the University of Wollongong Animal Ethics Committee (AEC) guidelines and regulations under protocol (AE16/10 and AE16/10r19) and SAHMRI’s Animal Ethics Committee. AEC guidelines comply with the ‘Australian code of practice for the care and use of animals for scientific purposes’, and the ARRIVE guidelines on reporting experiments involving animals (33). For University of Wollongong studies, adult male C57BL/6 mice (8-14 weeks old) were purchased from Australian Bioresources (Moss Vale, NSW, Australia) and housed in individually ventilated cages with a 12 h light/dark cycle; plastic shelter, nesting material, food pellets and water were available *ad libitum*. For studies performed at SAHMRI, male C57BL/6J mice aged 13–17 weeks were used. These mice were acquired from an in-house C57BL/6J breeding programme (strain no. 000664; originally purchased from The Jackson Laboratory, barn MP14) within SAHMRI’s specific and opportunistic pathogen-free animal care facility.

### Human DRG

Four L1 DRGs were collected by AnaBios (San Diego, CA, USA) from human adult organ donors (22.2 ± 2.08 years of age; male:female ratio 2:2). The samples were preserved in RNAlater (Thermo Fisher Scientific) and shipped to SAHMRI for processing (8, 9). The AnaBios NIH compliant ethics statement is available at: https://anabios.com/ethics-statement/.

### Quantitative-reverse-transcription-PCR (QRT-PCR)

Total RNA was extracted from human L1 DRG or mouse thoracolumbar (T10-L1) DRGs using PureLink RNA isolation kit (ThermoFisher Scientific). QRT-PCR was conducted with 20 ng RNA/well using the EXPRESS One-Step Superscript kit (ThermoFisher Scientific) and predesigned mouse or human-specific Taqman^®^ probes for GABA_B_R subunits (*Gabbr1*, Mm00433461_m1 or Hs00559488_m1, *Gabbr2*, Mm01352554_m1 or Hs01554996_m1), Ca_V_2.2 (*Cacna1b*, Mm01333678_m1 or Hs01053090_m1), Ca_V_2.3 (*Cacna1e*, Mm00494444_m1 or Hs00167789_m1), GIRK1-4 (*Kcnj3*, Mm00434618_m1 or Hs04334861_s1, *Kcnj6*, Mm01215650_m1 or Hs01040524_m1, *Kcnj9*, Mm00434621_m1 or Hs05018005_m1, *Kcnj5*, Mm01175829_m1 or Hs00168476_m1), *Gapdh* (Mm99999915_g1), *Ppia* (Mm02342430_g1 or Hs99999904_m1), and *Actb* (Mm00607939_s1 or Hs99999903_m1). mRNA was transcribed into cDNA using Supserscript IV kit (ThermoFisher Scientific). All samples were measured in duplicates. Raw data were analysed using DA2 software (ThermoFisher Scientific) and exported to Microsoft Excel to calculate relative mRNA quantities before transferring data into GraphPad for graphical representation. The comparative cycle threshold method was used to quantify the abundance of target transcripts relative to the reference genes (*Gapdh, Ppia*, and *Actb*).

### Single-cell RT-PCR

A total of 53 single subserosal colon-traced dissociated TL DRG neurons were isolated from one healthy mouse (C57Bl/6J; male; 16 weeks old) and collected using a micromanipulator at 40x magnification. During the picking process, a continuous slow flow of sterile, RNA/DNAse-free PBS was applied to minimize contamination with ambient RNA. After isolating a neuron, the glass capillary containing the cell was broken into a tube containing 10 μl lysis buffer with DNAse (TaqMan^®^ Gene Expression Cells-to-CT™ Kit; ThermoFisher Scientific). The entire sample was used for cDNA synthesis (SuperScript^®^ VILO™ cDNA Synthesis Kit, Thermo Fisher Scientific) and diluted 1:3 to measure target expression per cell using RT-PCR for 50 cycles. For each coverslip, a bath control was collected and analysed together with samples. Expression of *Tubb3* (Mm00727586_s1) served as a positive control to confirm neuronal identity. Cells were excluded from analysis if they lacked *Tubb3* expression or tested positive for *Gfap* (Mm01253033_m1), indicating contamination with glial cells. Out of the initial 53 cells, 12 were excluded based on these criteria. The frequency of the expression of a particular target was calculated from the remaining 41 neurons.

### Cell culture for patch clamp recordings of mouse DRGs

Mice were euthanised by isoflurane inhalation followed by rapid decapitation. The thoracic and lumbar DRG were exposed via laminectomy, harvested, and immediately transferred to ice cold (4°C) free of Ca^2+^- and Mg^2+^-free Hanks Buffered Saline Solution (HBSS). The DRGs were trimmed, removing central and peripheral nerve processes, and digested in HBSS containing collagenase type II (3 mg.ml^-1^; Worthington Biomedical Corp., Lakewood, NJ, USA) and dispase (4 mg.ml^-1^; GIBCO, Australia) at 37°C for 40 min. Following digestion, the ganglia were rinsed three to four times with 37°C/5% CO_2_-equilibrated F-12/GlutaMAX™ (Invitrogen) supplemented with 10% heat-inactivated FBS (GIBCO™, ThermoFisher Scientific) and 1% Pen/Strep. Cells were dispersed by mechanical trituration with progressively smaller fire-polished glass Pasteur pipettes. The resulting cell suspension was filtered through a 160 μm nylon mesh (Millipore Australia Pty Ltd, North Ryde, NSW, Australia) to remove any undigested material. Dissociated DRG neurons were plated onto 12 mm cover glass coated with poly-D-lysine (Sigma-Aldrich, Australia) and allowed to settle for ∼3 hr at 37°C, supplemented with fresh media, and incubated overnight before patch clamp recordings.

### Whole-cell current-clamp electrophysiology of mouse DRG neurons

Current clamp recordings of DRG neurons were carried out within 24 hr of dissociation in a extracellular (bath) solution containing (in mM): 140 NaCl, 4 KCl, 2 CaCl_2_, 1 MgCl_2_, 10 HEPES and 10 glucose, pH 7.4 adjusted with NaOH (∼320 mOsmol.kg^-1^). Fire polished borosilicate patch pipettes (World Precision Instruments) had resistances of 2-4 MΩ when filled with an internal solution containing (in mM): 130 KCl, 20 NaCl, 5 EGTA, 5 MgCl_2,_ 10 HEPES, 5 Mg-ATP, 0.2 Na_2_GTP, pH 7.2 adjusted with KOH (∼300 mOsmol.kg^-1^). Whole-cell configuration was achieved in voltage clamp mode before transitioning to current clamp mode. Depolarizing current steps of 500 ms duration at 0.2 Hz were applied to elicit action potential firing in small to medium diameter (<30 μm) DRG neurons with resting membrane potentials (RMP) more negative than −40 mV.

Modulation of high voltage-activated N-type (Ca_V_2.2) channels by ω-conotoxin CVIE, and GIRK channels by the antagonist Tertiapin-Q (TPQ) and the agonist ML297, was conducted via bath application of these compounds. In a series of experiments the GABA_B_R agonist, α-conotoxin Vc1.1, was co-applied with either CVIE or ML297. All drugs were dissolved in distilled H_2_O to prepare to stock solutions at the appropriate concentration. The extracellular solutions were superfused using a peristaltic pump at a flow rate of 1 ml min^−1^ in a ∼500 µL experimental chamber maintained at room temperature (21-23°C). TPQ was purchased from Abcam (Cambridge, UK), ML297 was obtained from Tocris (Bristol, UK), and ω-conotoxin CVIE and α-conotoxin Vc1.1 were synthesized as described previously (Berecki et al., 2010) and kindly provided by Profs David Craik and Richard Clark (The University of Queensland, Brisbane, Australia).

### Mouse model of chronic visceral hypersensitivity (CVH)

Colitis was induced by the administration of 2,4-dinitrobenzene sulfonic acid (DNBS), as described previously (34). Briefly, 13–15-week-old anaesthetized mice were administered a single intra-colonic enema of 100 µL DNBS/Ethanol solution (5 mg DNBS in 100 µL of 30% ethanol) via a polyethylene catheter to induce colonic inflammation (9, 35). Histological examination of mucosal architecture, cellular infiltrate, crypt abscesses, and goblet cell depletion confirmed significant DNBS-induced damage by day 3 post-treatment. This damage largely resolved spontaneously by day 7 and was fully resolved by day 28 (36). High-threshold colonic nociceptors from mice at the 28-day time point displayed significant mechanical hypersensitivity and reduced mechanical activation thresholds (8, 9, 13, 14, 29, 34–39). This model is characterized by hyperalgesia and allodynia to colorectal distension and is therefore termed ‘Chronic Visceral Hypersensitivity’ (CVH) (9, 13, 14, 29, 34, 38, 40).

### Retrograde tracing to identify colon-innervating DRG neurons

CTB-conjugated to AlexaFluor^®^-488 or −555 (ThermoFisher Scientific) was injected (4 μL/injection) subserosally at three sites within the distal colon of healthy mice (single cell RT-PCR and patch clamp recordings) or CVH mice (patch clamp recordings) using a Hamilton syringe fitted with a 23-gauge needle as described previously (41).

### Cell culture for patch clamp recordings and single-cell picking of colonic DRGs

Four days after retrograde tracing, mice were humanely euthanized, and thoracolumbar (TL, from T9–L1) DRG were removed and dissociated (8, 9). DRGs were enzymatically digested first with 4 mg ml^−1^ collagenase II GIBCO, ThermoFisher Scientific) and 5.3 mg ml^−1^ dispase I (GIBCO, ThermoFisher Scientific) for 30 min at 37 °C, followed by a second digestion with 4 mg ml^−1^ collagenase II for 10 min at 37 °C. The dissociated neurons were resuspended in DMEM (GIBCO, ThermoFisher Scientific) supplemented with 10% fetal calf serum (Invitrogen, ThermoFisher Scientific), 2 mM L-glutamine (GIBCO, ThermoFisher Scientific), 100 μM MEM non-essential amino acids (GIBCO, ThermoFisher Scientific), 96 ng ml^−1^ NGF, and 100 mg ml^−1^ penicillin/streptomycin (GIBCO, ThermoFisher Scientific). Neurons were spot-plated onto coverslips coated with laminin (20 μg ml^−1^; Sigma-Aldrich) and poly-D-lysine (800 μg ml^−1^; ThermoFisher Scientific) and maintained in an incubator at 37 °C with 5% CO_2_.

### Whole-cell current-clamp electrophysiology of colon-innervating DRG neurons

Dissociated DRG neurons isolated from colon-traced healthy or CVH mice were recorded on day 1 post-culture (20-30 hs after plating) (14, 29, 38). The intracellular current-clamp solution contained (in mM): 135 KCl; 2 MgCl_2_; 2 MgATP; 5 EGTA-Na; 10 HEPES-Na; adjusted to pH 7.3. The extracellular (bath) current-clamp solution contained (in mM): 140 NaCl; 4 KCl; 2 MgCl_2_; 2 CaCl_2_; 10 HEPES; 5 glucose; adjusted to pH 7.4. Standard-wall borosilicate glass pipettes (OD x ID x length: 1.5 mm x 0.86 mm x 7.5 cm, Harvard) pulled and fire-polished to a resistance of 3 - 6 MΩ using a P-97 (Sutter Instruments) pipette puller were used. The protocol used was as follows, in current-clamp mode, neurons were held at −70 mV for 15 ms, hyperpolarized with a −20 pA current injection for 475 ms, then returned to −70 mV for 100 ms. Stepwise depolarizing pulses were applied in 0 pA or 25 mV increments (475 ms) from a holding potential of −70 mV, with 2 sec repetition intervals, to determine the rheobase (minimum amount of current required to elicit an action potential). Rheobase was assessed both in the presence and absence of Tertiapin-Q (100 nM) or the HCN inhibitor ZD7288 (50 μM). Neurons with a resting membrane potential more depolarized than −40 mV were excluded from recordings, as this indicated poor cell health. Recordings were amplified using an Axopatch 200A, digitized with a Digidata 1322A, sampled at 20 kHz, filtered at 5 kHz, and recorded with pCLAMP 9 software (Molecular Devices). Data was analyzed using Clampfit 10.3.2 (Molecular Devices) and Prism v9.3.1 (GraphPad).

### *Ex vivo* electrophysiology

Recordings were obtained from healthy male C57BL/6J mice. Pelvic and splanchnic afferent nerve activity was measured *ex vivo* using an isolated segment of the mouse colon ligated at either end to allow for fluid distension (42). The tissue was mounted in a custom-built organ bath and perfused with warm, carbogenated physiological Krebs buffer (composition in mmol/L: 118.4 NaCl; 24.9 NaHCO_3_; 1.9 CaCl_2_; 1.2 MgSO_4_; 4.7 KCl; 1.2 KH_2_PO_4_; 11.7 glucose). To suppress smooth muscle activity, the Krebs solution included the L-type calcium channel antagonist nifedipine (1 µM). The prostaglandin synthesis inhibitor indomethacin (3 µM) was also added to suppress potential inhibitory actions of endogenous prostaglandins. The pelvic and splanchnic nerve were carefully isolated and placed within a sealed glass pipette containing a microelectrode (WPI) connected to a NeuroLog headstage (NL100AK; Digitimer). Nerve activity was amplified (NL104), filtered (NL 125/126, bandpass 50–5,000 Hz, NeuroLog; Digitimer), and digitised using a CED 1401 interface (Cambridge Electronic Design, Cambridge, UK) for offline analysis with Spike2 software (Cambridge Electronic Design, Cambridge, UK). Nerve activity was measured at baseline and during ramp distension with Krebs solution or Tertiapin-Q (flow rate: 100 µl/min; pressure range: 0 to 80 mmHg) measured with a pressure transducer; NL108T2 Digitimer).

The number of action potentials exceeding a preset threshold (twice the background electrical noise) was determined per second to quantify the afferent response. Responses were compared before and after the intraluminal application of Tertiapin-Q (10 μM and 100 μM). Single-unit analysis of action potentials was conducted offline by matching individual spike waveforms through linear interpolation using Spike2 v.10.05 software (Cambridge Electronic Design, Cambridge, UK).

### Statistical analysis

Data are expressed as mean ± SEM or as the percentage of neurons. Figures were prepared using GraphPad Prism 9 software (San Diego, CA, USA). *N* represents the number of animals, and *n* denotes the number of neurons or afferents. Statistical significance was defined as *P*□<□0.05 (\**), P*□*<*□*0.01* (**)*, P*□*<*□*0.001* (***), and *P*□<□0.0001 (****).

All data were analysed using GraphPad Prism 9 and assessed for normal distribution using either the Kolmogorov–Smirnov or Shapiro–Wilk test. Depending on the experimental design, data were further analyzed using one-way ANOVA followed by Tukey’s post hoc test for multiple group comparisons, two-way ANOVA followed by Bonferroni’s post hoc test, or paired/unpaired two-tailed *t*-tests. The specific statistical tests used for each dataset are detailed in the corresponding figure legends.

## RESULTS

### Expression of GABA_B_R subunits, Ca_V_2.2, Ca_V_2.3, and GIRK1-4 in mouse and human thoracolumbar DRGs

Quantitative RT-PCR (QRT-PCR) analysis of whole mouse DRG samples revealed that GABA_B_R1, Ca_V_2.2 and GIRK1 were the most abundant targets expressed in the thoracolumbar dorsal root ganglia (DRGs, T10-L1) followed by Ca_V_2.3 and GABA_B_R2. In contrast, GIRK2-4 exhibited very low mRNA expression levels, 51-times lower than GIRK1 for GIRK2, 106-times lower for GIRK4, and 306-times lower for GIRK3 (**Fig. 1A-B**). In mouse DRGs, the fold change in expression for each target relative to the least expressed gene was as follows: GABA_B_R1 (1961) > Ca_V_2.2 (1188) > GIRK1 (306) > Cav2.3 (66) > GABA_B_R2 (34) > GIRK2 (6) > GIRK4 (3) > GIRK3 (1). A similar expression pattern was observed in human thoracolumbar DRGs (L1, **Fig. 1C-D**). However, GABA_B_R2 expression was relatively higher in human DRGs, being four-fold lower than GABA_B_R1, compared to mouse DRGs, where GABA_B_R2 was 60-fold lower than GABA_B_R1. Additionally, the expression of Cav2.3 in human DRGs was notably lower than in mice, with Ca_V_2.3 being the least expressed target in human DRGs. Similar to mouse TL DRG, Cav2.2 was highly expressed in human L1 DRG, however, the ratio of GABA_B_R1 to Ca_V_2.2 was only 12 to 1 compared to the similar expression levels in mice (1.6 to 1). In human DRGs, the fold change in expression for each target relative to the least expressed gene was as follows: GABA_B_R1 (1231) > GABA_B_R2 (365) > GIRK1 (163) > Ca_V_2.2 (99) > GIRK3 (13) > GIRK2 (5) > GIRK4 (3) > Ca_V_ 2.3 (1). Overall, these findings highlight that Ca_V_2.2, GABA_B_R1, and GIRK1 are the dominant isoforms in both mouse and human DRGs, suggesting their potential functional importance in thoracolumbar sensory processing.

**Figure 1.**
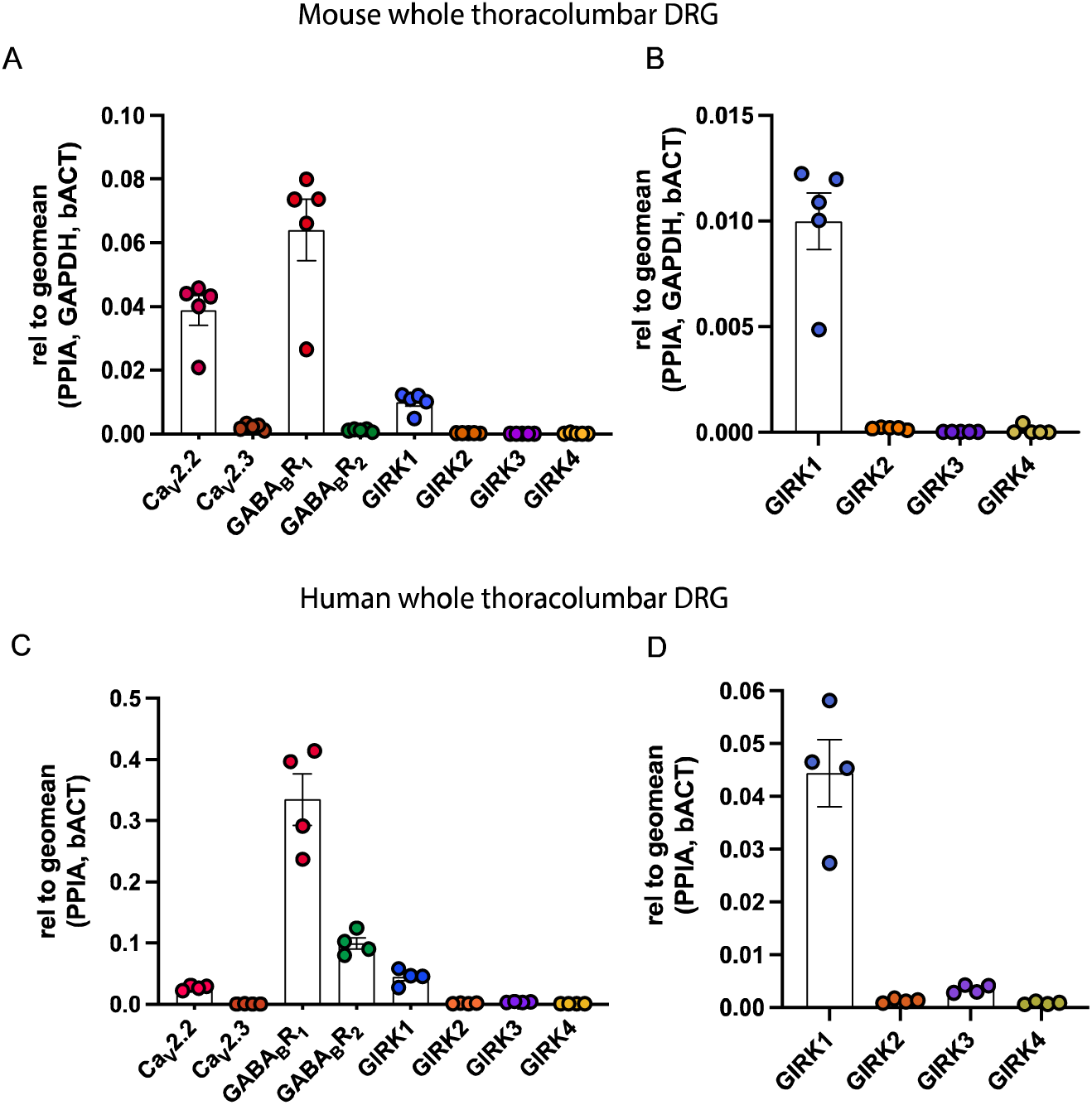
Expression patterns of GABA_B_ receptors and associated ion channels in mouse and human thoracolumbar DRG (T10-L1). **(A)** Quantitative RT-PCR (QRT-PCR) analysis showing high mRNA abundance of GABA_B_R1, Ca_V_2.2, and GIRK1 in whole mouse thoracolumbar DRG. Data represent 5 individual mice, with each dot representing one mouse. **(B)** Inset of (A), showing QRT-PCR analysis of GIRK1-4 mRNA expression in whole mouse thoracolumbar DRG (T10-L1). Results indicate low mRNA abundance for GIRK2-4. **(C)** QRT-PCR analysis of mRNA expression in whole human thoracolumbar (L1). Results show high expression of GABA_B_R1, GABA_B_R2, and GIRK1, while Ca_V_2.2 is expressed at relatively lower levels compared to the abundant GABA_B_R1 expression. **(D)** Inset of (C), showing QRT-PCR analysis of GIRK1-4 expression in human L1 DRG, indicating low mRNA abundance of GIRK2-4. Data are from DRG samples of 4 human donors, with each point representing an individual donor.

### Co-expression of GABA_B_R subunits, Ca_V_2.2, Ca_V_2.3, and GIRK channels in colon-innervating DRGs

Single-cell RT-PCR from retrogradely traced mouse colon-innervating DRGs was performed to assess the expression of GABA_B_R, Ca_V_ channels, and GIRK channels specifically in colonic DRG neurons. The analysis revealed high expression levels of Ca_V_2.2 and GABA_B_R2, detected in 100% and 95% of colon-innervating DRG neurons, respectively (**Fig. 2A-B**). In contrast, Ca_V_2.3 was expressed in 29 of 41 cells (71%), and GABA_B_R1 was expressed in 26 of 41 neurons (63%, **Fig. 2A-B**). Among the 41 colonic neurons analyzed, GIRK1 was expressed in 24 neurons (58%), whereas GIRK2 was detected in only 6 neurons (15%). GIRK3 and GIRK4 expression levels were below the detection threshold and were not observed in any of the neurons analyzed. Co-expression analysis revealed that when GIRK1 was expressed it was highly co-expressed with Ca_V_2.2 (100%), Ca_V_2.3 (75%), GABA_B_R1 (71%), and GABA_B_R2 (96%) (**Fig. 2C**), suggesting a strong functional link between GABA_B_R signaling, Ca_V_ channels, and GIRK channel activity in colonic sensory DRG neurons.

**Figure 2.**
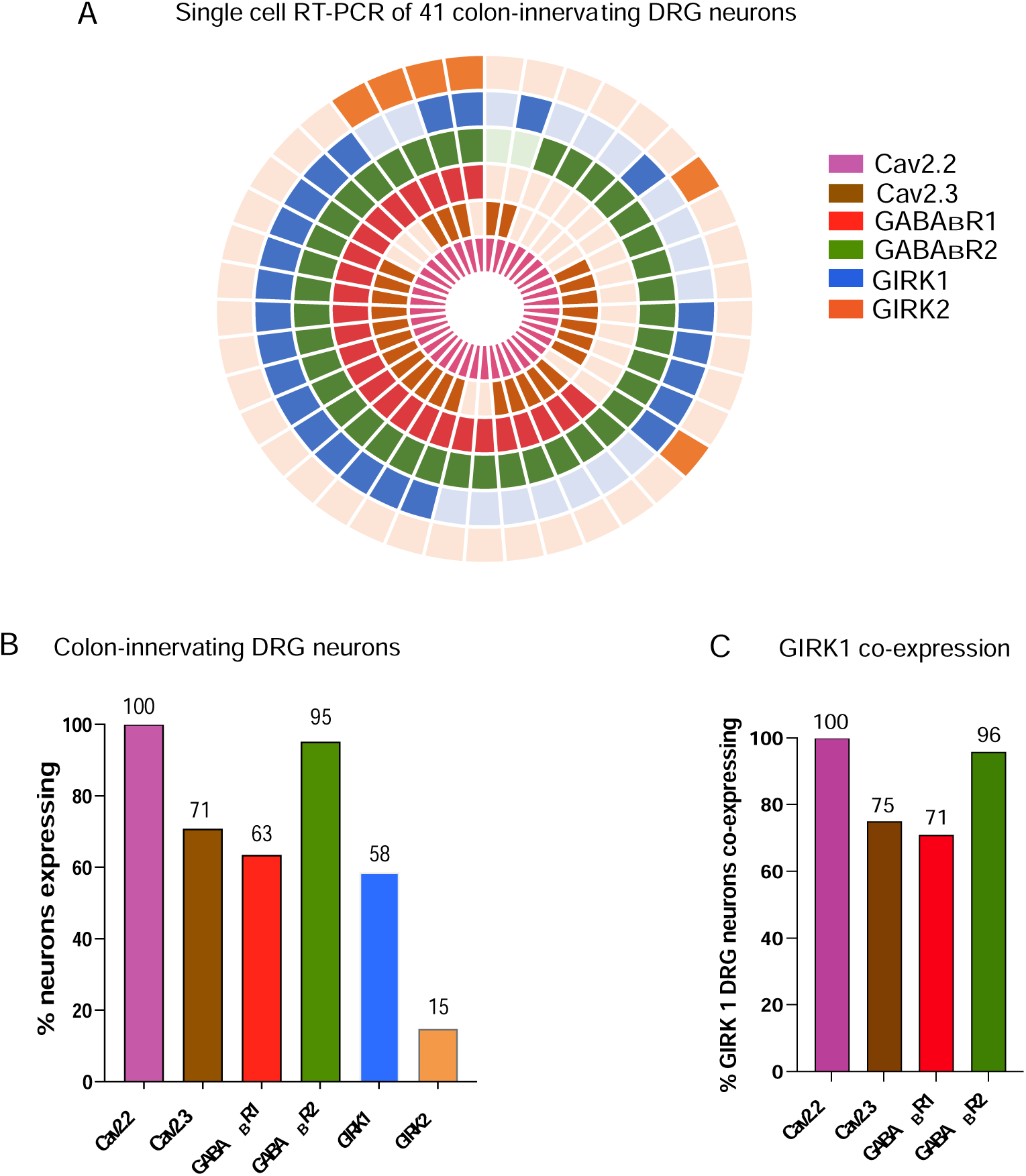
Expression and co-expression of ion channel and receptor genes in retrogradely traced colon-innervating mouse DRG neurons. **(A)** Donut plot illustrating the expression and co-expression of genes encoding Ca_V_2.2, Ca_V_2.3, GABA_B_R1, GABA_B_R2, and GIRK1-2 in 41 individual retrogradely traced colon-innervating DRG neurons (N = 1 mouse). Each gene is represented by a distinct colour, with expression indicated by bold shading. GIRK2 is depicted in the outer ring, while the inner ring represents Ca_V_2.2. Individual neurons are arranged radially, allowing for clear visualisation of gene co-expression within a single neuron from the outermost to the innermost rings. Some neurons expressed most targets, while others showed combinations of specific targets. (**B**) Single-cell RT-PCR analysis of retrogradely traced colon-innervating DRG neurons showing the percentage of colon-innervating DRG neurons expressing each transcript: Ca_V_2.2 (100%), Ca_V_2.3 (71%), GABA_B_R1 (63%), GABA_B_R2 (95%), GIRK1 (58%), and GIRK2 (15%). No expression of GIRK3 or GIRK4 was detected. **(C)** Single-cell RT-PCR analysis highlighting the gene co-expression of GIRK1 with Ca_V_2.2 (100%), Ca_V_2.3 (75%), GABA_B_R1 (71%), and GABA_B_R2 (96%).

### Effects of the selective Ca_V_2.2 antagonist **ω**-conotoxin CVIE and the GIRK channel agonist ML297 on mouse DRG neuronal excitability

To investigate the roles of Ca_V_2.2 and GIRK channels in regulating neuronal excitability, whole-cell patch clamp recordings were performed on mouse DRG neurons in the presence of the selective Ca_V_2.2 antagonist ω-conotoxin CVIE and the GIRK channel agonist ML297 (**Fig. 3**). Application of ω-conotoxin CVIE (100 nM) significantly reduced action potential (AP) firing frequency by approximately 50% and increased rheobase by approximately 20%, without affecting the resting membrane potential (−64.5 ± 2.9 mV, n = 6, **Fig. 3A-B**). Co-application of ω-conotoxin CVIE with α-conotoxin Vc1.1 caused membrane hyperpolarization and a further reduction in action potential (AP) firing (**Fig. 3A**). Together, co-application of ω-conotoxin CVIE and α-conotoxin Vc1.1 significantly decreased the resting membrane potential and input resistance, reduced AP firing frequency, and significantly increased rheobase (**Fig. 3B**), indicating a synergistic suppression of excitability in DRG neurons.

**Figure 3.**
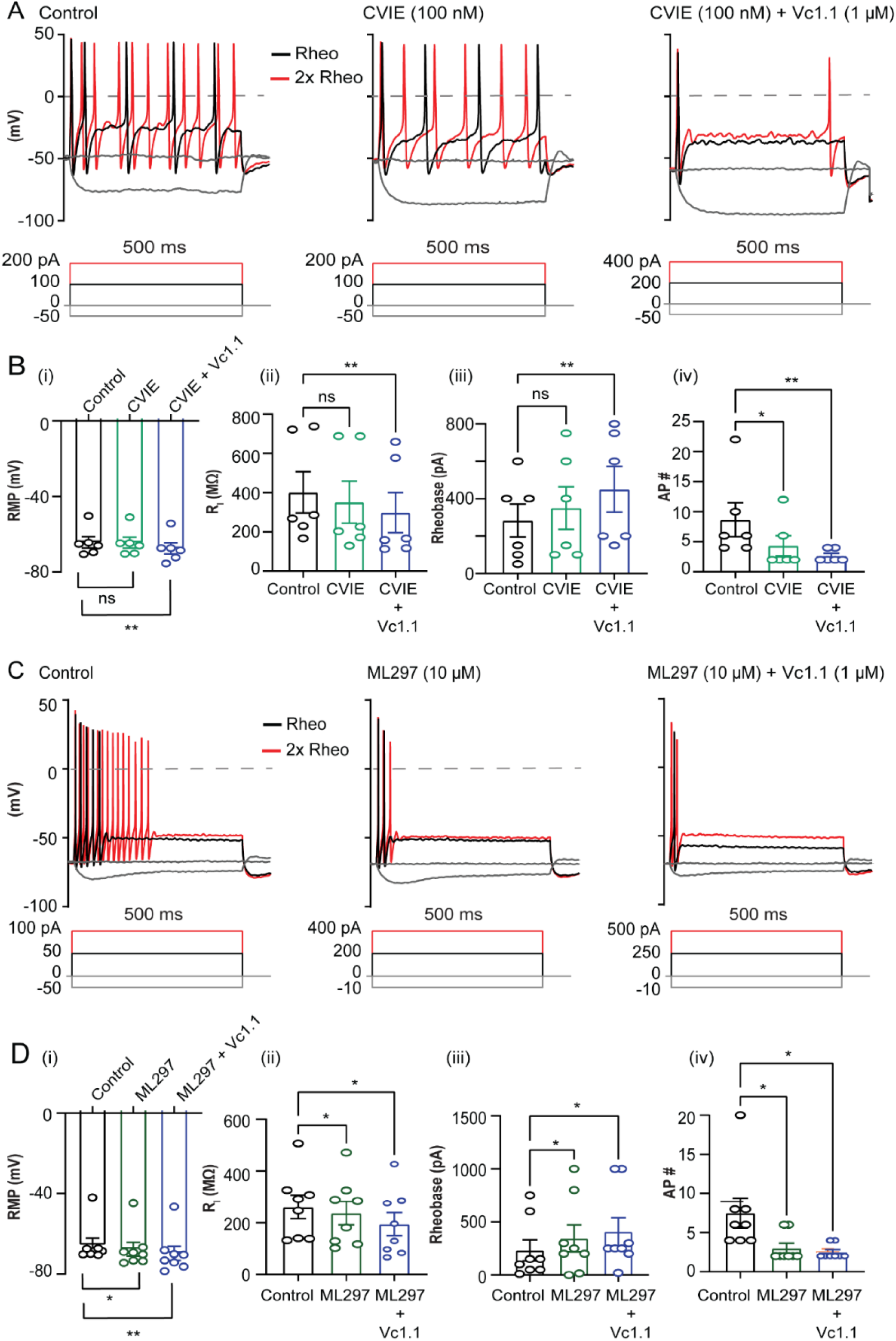
Effects of ω-conotoxin CVIE, ML297, and α-conotoxin Vc1.1 on the electrophysiological properties of adult mouse DRG neurons. **(A)** Representative voltage responses to current clamp steps recorded in a small diameter DRG neuron (23 μm) in the absence (control; **left**), and presence of CVIE (100 nM) (**middle**), and CVIE (100 nM) + Vc1.1 (1 µM) (**right**). The dashed line indicates 0 mV. Rheobase (black) and 2x rheobase (red) are shown for both membrane potential and current. (**B**) Bar graphs summarizing the effects of CVIE and CVIE + Vc1.1 on (**i**) resting membrane potential (RMP), (**ii**) input resistance (Ri), (**iii**) rheobase, and (**iv**) AP frequency (Hz) in response to 500 ms depolarizing current steps in small to medium diameter DRG neurons from adult mice. (**C**) Representative voltage responses to current clamp steps recorded in a mouse DRG neuron (24.5 μm) in the absence (control; **left**) and presence of ML297 (10 µM) (**middle**), and ML297 (10 µM) + Vc1.1 (1 µM) (**right**). The dashed line denotes 0 mV. Rheobase (black) and 2x rheobase (red) are shown for both membrane potential and current. (**D**) Bar graphs summarizing the effects ML297 and ML297 + Vc1.1 on (**i**) resting membrane potential (RMP), (**ii**) input resistance (Ri), (**iii**) rheobase, and (**iv**) AP frequency (Hz) in response to a 500 ms depolarizing current step at 2x rheobase in adult mouse DRG neurons (< 30 μm diameter). Data are presented as mean ± SEM, with the number of neurons indicated within each bar (n = 8). Statistical significance was determined using one-way repeated measures ANOVA or Friedman test, * P < 0.05.

Similarly, application of the GIRK channel agonist ML297 (10 µM) produced effects comparable to ω-conotoxin CVIE on resting membrane potential, rheobase, input resistance, and AP firing frequency (**Fig. 3C**). However, unlike ω-CVIE, ML297 caused significant membrane hyperpolarization (>3.5 mV, n = 8). These findings are consistent with previous studies showing that GABA_B_R-dependent GIRK channel potentiation by baclofen or Vc1.1 causes membrane hyperpolarization and reduced excitability in mouse DRG neurons (32). Furthermore, co-application of ML297 and Vc1.1 (1 μM) produced an additive effect, further increasing the current required to induce to trigger AP firing and significantly reducing firing frequency in healthy mouse DRG neurons (**Fig. 3C-D**, **Table 1**).

**Table 1.** Passive and active electrical properties of adult mouse DRG neurons in the absence (Control) and presence of either ω-conotoxin CVIE (100 nM) or ML297 (10 μM) applied alone or in combination with α-conotoxin Vc1.1 (1 µM). Parameters analysed include resting membrane potential (RMP, mV), input resistance (Ri, MΩ), rheobase (pA), and action potential (AP) frequency (Hz) in response to a 500 ms depolarizing current step. Results are presented as mean ± SD, with the number of experiments indicated. Statistical significance was determined using one-way repeated measures ANOVA or the Friedman test (*P < 0.05).

| | Control | ω-CVIE | ω-CVIE<br>+Vc1.1 | n | Control | ML297 | ML297+Vc1.1 | n |
| --- | --- | --- | --- | --- | --- | --- | --- | --- |
| Rheobase (pA) | 283.3 ± 88.2 | 350 ± 14.0 | 450 ± 122.5 | 6 | 232.5 ± 99.1 | 358.1 ± 123.7 | 408.8 ± 132.4 | 8 |
| Input Resistance (MΩ) | 401.7 ± 105.0 | 351.4 ± 107.3 | 298.4 ± 101.9 | 6 | 260.9 ± 42.1 | 237.3 ± 45.2 | 195.0 ± 44.5 | 8 |
| RMP (mV) | -64.3 ± 2.9 | -64.5 ± 2.9 | -43.4 ± 23.3 | 6 | -65.6 ± 3.4 | -68.8 ± 3.6 | -69.7 ± 3.5 | 8 |
| AP (Hz) | 8.7 ± 2.8 | 4.3 ± 1.7 | 2.7 ± 0.4 | 6 | 7.5 ± 1.9 | 3.0 ± 0.7 | 2.5 ± 0.3 | 8 |

### GIRK channel inhibitor Tertiapin-Q increases the excitability of adult mouse DRG neurons, and its effects are counteracted by **α**-conotoxin Vc1.1

To further examine the role of GIRK channels in regulating neuronal excitability, we applied the specific GIRK channel antagonist Tertiapin-Q (TPQ; 100 nM) to adult mouse DRG neurons. Application of TPQ caused membrane depolarization and a significant increase in neuronal excitability (**Fig. 4A-B**, **Table 1**). Specifically, TPQ significantly elevated the resting membrane potential and input resistance, increased action potential firing frequency, and reduced rheobase. To determine whether TPQ could interfere with GIRK channel potentiation by α-conotoxin Vc1.1, we co-applied TPQ and Vc1.1. Co-incubation increased AP discharge relative to Vc1.1 alone, indicating that TPQ blocked the potentiation of GIRK channel currents induced by Vc1.1. Under control conditions, Vc1.1 potentiates GIRK channel activity, leading reduced AP firing. Thus, the excitatory effects observed with TPQ in the presence of Vc1.1 suggest that GIRK channel inhibition reverses the Vc1.1-induced suppression of neuronal excitability **(Fig. 4C-D)**.

**Figure 4.**
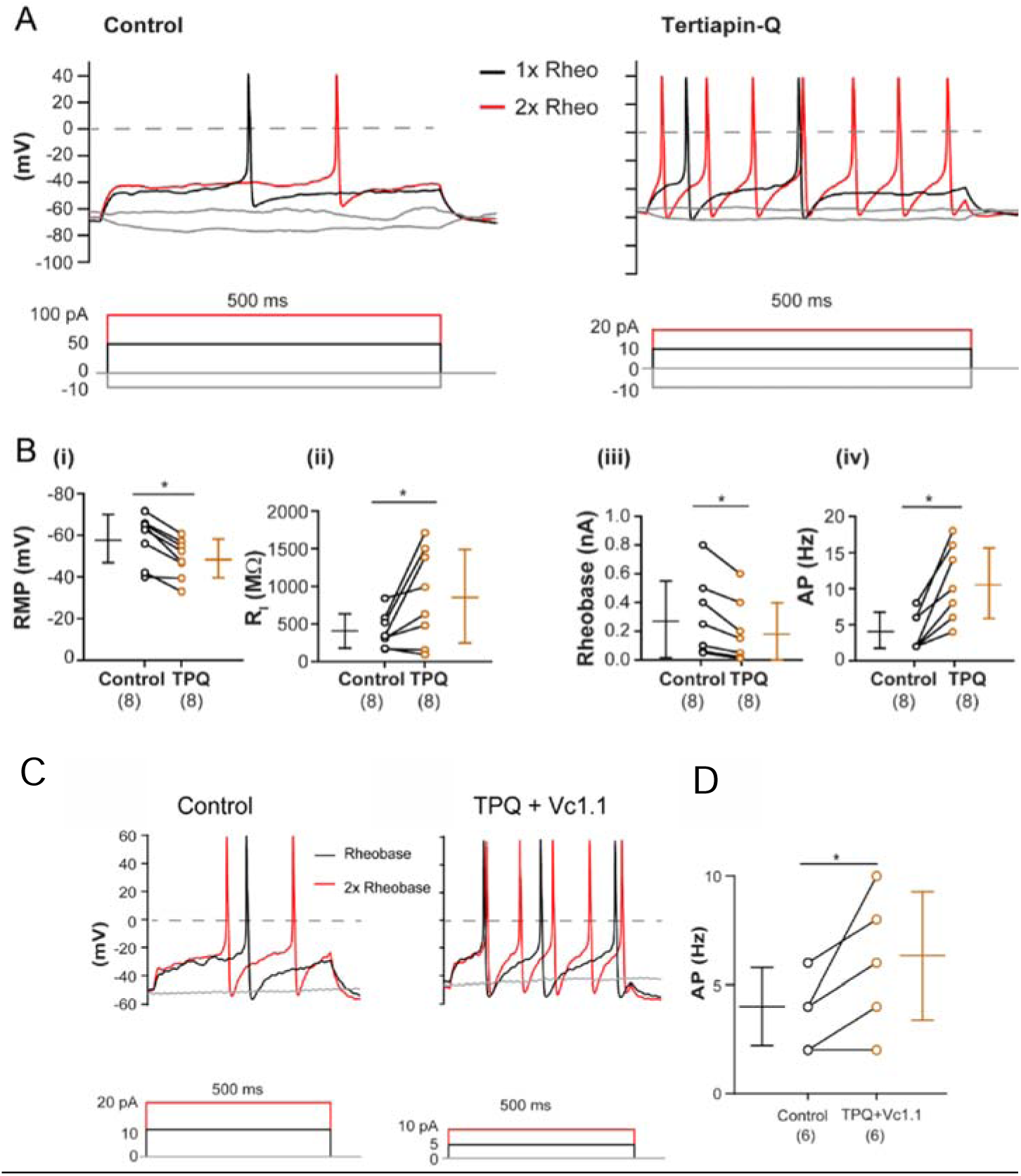
The GIRK channel inhibitor Tertiapin-Q (TPQ) enhances the excitability of adult mouse DRG neurons. **(A)** Representative voltage responses to current clamp steps recorded in a mouse DRG neuron (26 μm diameter) in the absence (control; **left**) and presence of 100 nM TPQ (**right**). The dashed line denotes 0 mV. Rheobase (black) and 2x rheobase (red) are shown for both membrane potential and injected current. **(B)** Paired scatter dot plots illustrating the effects TPQ (100 nM) on (**i**) resting membrane potential (RMP), (**ii**) input resistance (R_i_), (**iii**) rheobase (nA), and (**iv**) AP frequency (Hz) in response to a 500 ms depolarizing current step at 2x rheobase in small to medium diameter (<30 μm) adult mouse DRG neurons. **(C, D)** Lack of effect of α-conotoxin Vc1.1 on neuronal excitability in the presence of Tertiapin-Q (TPQ). Representative voltage responses to current clamp steps recorded in a small to medium diameter mouse DRG neuron recorded in the absence (control) and presence of 100 nM TPQ + 1 µM α-conotoxin Vc1.1 **(C)**. Insets show responses to 500 ms depolarizing current steps. The dashed line denotes 0 mV, while rheobase (black) and 2x rheobase (red) are indicated for both membrane potential and current. **(C)** Paired scatter dot plot showing action potential (AP) frequency (Hz) at 2x rheobase under control conditions and in the presence of TPQ (100 nM) + Vc1.1 (1 µM) in small to medium diameter (<30 μm) adult mouse DRG neurons. Data are presented as mean ± SD, with the number of neurons indicated in parentheses. Statistical analysis was performed using a paired t-test or the Wilcoxon matched-pairs signed rank test; * P < 0.05.

### GIRK channel inhibitor Tertiapin-Q does not affect the responsiveness of mouse colonic afferents

To evaluate the functional role of GIRK channels in colonic sensory pathways, we conducted *ex vivo* recordings of pelvic and splanchnic nerve activity in response to colonic distention. Afferent recordings from pelvic nerves revealed that as intraluminal pressure increased, action potential firing also increased proportionately **(Fig. 5A)**. However, comparison of responses during colonic distension in the presence of vehicle (Krebs solution) versus intraluminal infusion of TPQ (10 μM or 100 μM) showed no significant differences in afferent firing rates or colonic compliance (**Fig. 5A-D)**. These findings indicate that GIRK channel inhibition by TPQ does not affect the excitability of healthy colonic pelvic afferents responding to mechanical distension. A more detailed analysis of individual afferent responses was performed by classifying pelvic colonic afferents based on their firing characteristics at specific distension pressures (42). TPQ treatment did not alter the mechanical sensitivity of low-threshold (LT), wide dynamic range (WDR), or high-threshold (HT) pelvic afferents (**Fig. 5E**), consistent with the whole pelvic nerve recordings.

**Figure 5.**
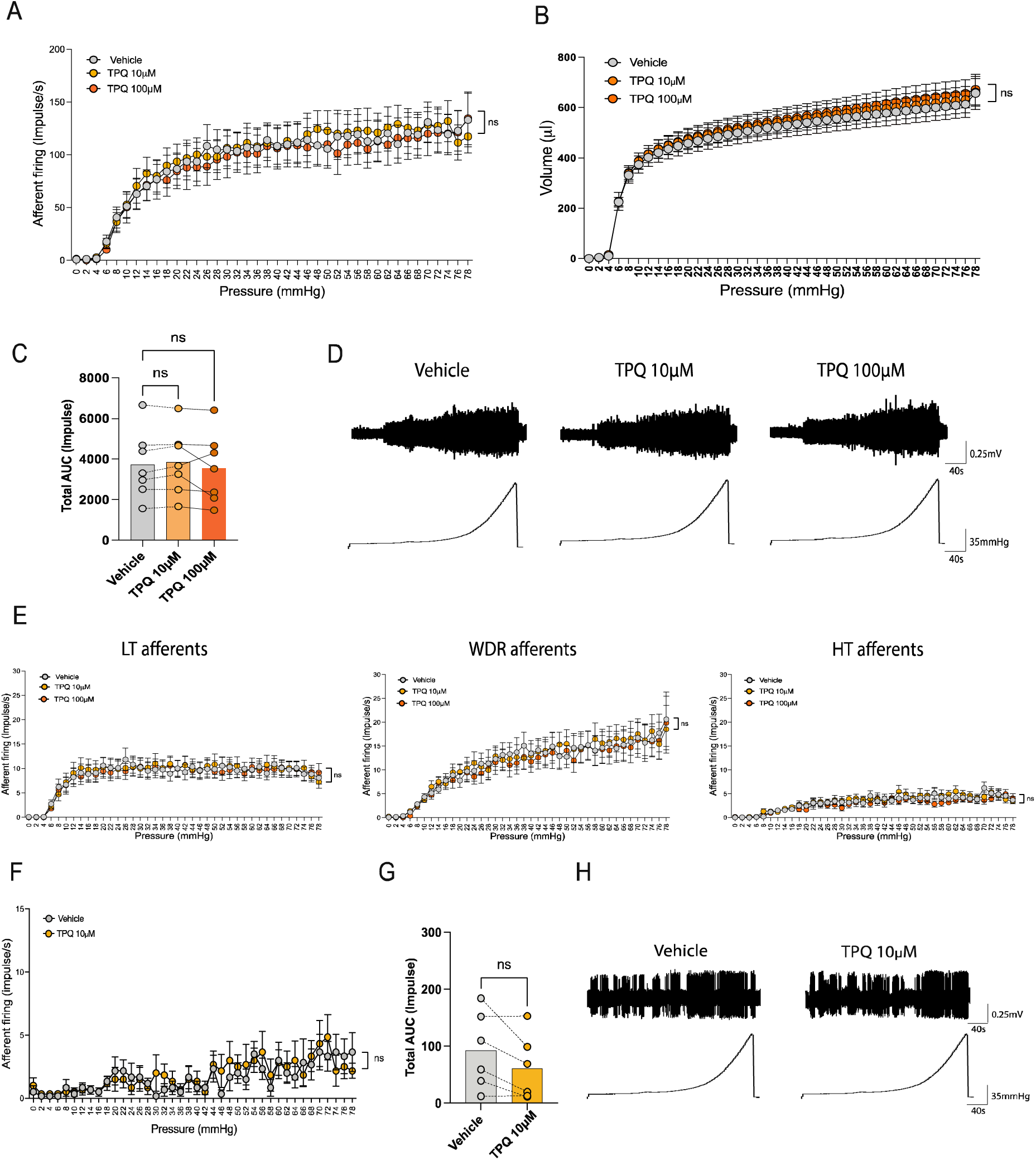
Lack of effect of Tertiapin-Q (TPQ) on pelvic and splanchnic afferent responses to colonic distension. **(A)** Group data showing pelvic nerve afferent firing responses and colonic compliance **(B)** to increasing distension pressures in *ex vivo* preparations from healthy C57BL/6J mice (N = 7). Responses were recorded at baseline (vehicle-Krebs distension) and following intraluminal application of TPQ at 10 μM or 100 μM. **(C)** Comparison of the total AUC for afferent firing between vehicle and TPQ intraluminal application, showing no significant differences (ns, P > 0.05). **(D)** Representative traces of pelvic nerve afferent firing in response to graded colonic distension in the absence (Vehicle, left) and following TPQ application (10 and 100 μM, right). **(E)** Post-hoc single-unit analysis of multiunit *ex vivo* colonic recordings. Pelvic nerve afferents were classified based on their discharge profile during ramp distension into low-threshold (LT), wide dynamic range (WDR), or high-threshold (HT). Total AUC for LT (n = 33), WDR (n = 25), and HT (n = 24) afferents during graded distension of the colon showed no significant effect of TPQ (10 μM or 100 μM). **(F)** Group data showing splanchnic nerve afferent firing responses to increasing colonic distension pressures in *ex vivo* preparations from healthy C57BL/6J mice (N = 6). Responses were recorded under baseline conditions (vehicle-treated Krebs distension) and following intraluminal application of TPQ (10 μM). **(G)** Comparison of total AUC for splanchnic afferent firing between vehicle-treated and TPQ-treated conditions, showing no significant differences (ns, P > 0.05). **(H)** Representative traces of splanchnic nerve afferent firing in response to colonic distension in the absence (Vehicle, left) and presence of TPQ (10 μM, right). Data are presented as mean ± SEM. P values are based on two-way ANOVA (Šidak’s multiple-comparisons test) or one-way ANOVA (Friedman’s multiple-comparisons test) as appropriate. N = 7 mice.

We also examined the effects of TPQ on splanchnic afferent responses to colonic distention (**Fig. 5F-H**). A small, non-significant decrease in splanchnic afferent firing was observed following intracolonic infusion of TPQ (10 μM) compared to vehicle (Krebs solution, **Fig. 5F-G**). Given that the majority of splanchnic afferents are high threshold, no further analysis of afferent subtypes was performed. Taken together, these results indicate that inhibiting GIRK channel activity with TPQ does not significantly affect the excitability of either pelvic or splanchnic colonic afferents in healthy mice during mechanical distension.

### GIRK channel inhibitor Tertiapin-Q does not affect the excitability of colon-innervating DRG neurons in CVH states

To further assess the role of GIRK channels on the excitability of colonic DRG neurons, we performed whole-cell patch clamp recordings in our CVH model. Recordings were obtained from retrogradely traced colonic DRG neurons isolated from CVH mice in the presence of the GIRK channel inhibitor Tertiapin-Q (TPQ, 100 nM). In contrast to our findings in general DRG neurons, TPQ application did not significantly alter the amount of injected current required to evoke an action potential, indicating no change in rheobase (**Fig. 6A-B**). Furthermore, TPQ had no significant effect on the resting membrane potential of colonic DRG neurons from CVH mice (**Fig. 6C**). These findings suggest that inhibiting GIRK channels with TPQ does not influence action potential firing in colon-innervating DRGs under CVH conditions. Taken together with our findings in healthy tissue, this indicates that GIRK channels may not play a prominent role in modulating colonic neuronal excitability in either physiological or pathophysiological states.

**Figure 6.**
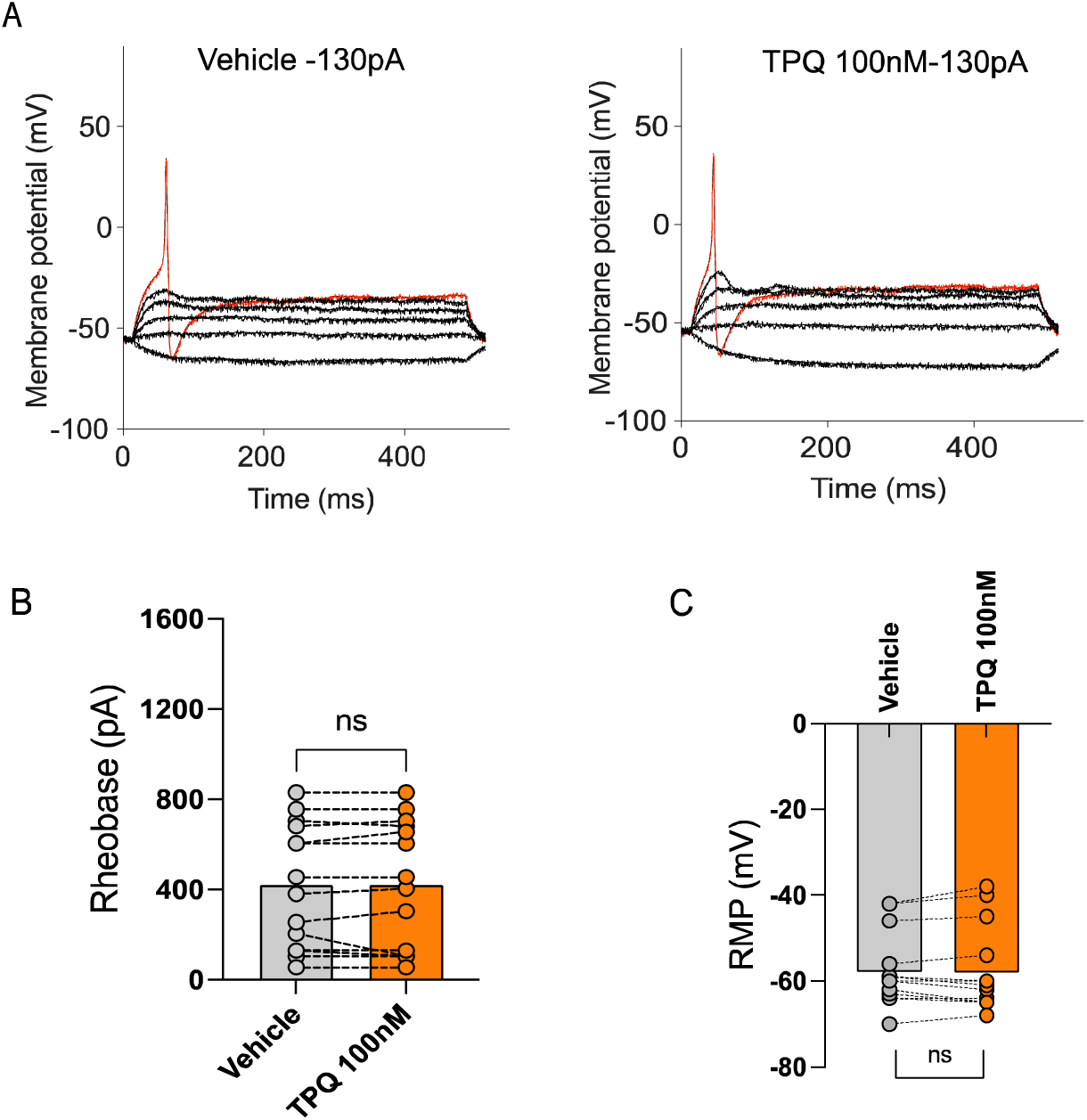
Lack of effect of Tertiapin-Q (TPQ) on the excitability of mouse colon-innervating DRG neurons in CVH states. **(A)** Representative voltage responses to depolarizing current clamp steps recorded from a thoracolumbar (TL) DRG neuron in the absence (control) and presence of TPQ (100 nM). Rheobase is indicated in red (130 pA). **(B, C)** Bar graphs showing no significant effect of TPQ (100 nM) on rheobase or resting membrane potential (RMP) in colon-innervating DRG neurons. Data are presented as mean ± SEM. *P* values are based on paired t-tests. n = 14 neurons from N = 3 mice.

### The specific HCN channel inhibitor ZD7288 increases the excitability of healthy colonic DRG neurons

A recent study in hPSC-derived sensory neurons demonstrated that although GIRK channels regulate neuronal excitability, their modulation by GABA_B_Rs was not observed, suggesting that functional GIRK1/2 channels were not coupled to GABA_B_Rs in these cells (31). However, this study identified a role for hyperpolarisation-activated cyclic nucleotide-gated (HCN) channels in regulating neuronal excitability. Specifically, application of the HCN inhibitor ZD7288 (30 μM) significantly altered both passive and active membrane properties in hPSC-derived sensory neurons (31). Given these findings, and our own observation that the GIRK inhibitor Tertiapin-Q had no significant effect on excitability in colonic DRG neurons and afferents, we investigated whether HCN channels contribute to the regulation of excitability in mouse colonic DRG neurons. To test this, we applied 50 μM ZD7288 and assessed the electrophysiological properties of healthy mouse colonic DRG neurons using whole-cell patch clamp recordings. Application of ZD7288 significantly increased the neuronal excitability, as evidenced by a significant decrease in the amount of injected current required to evoke an action potential (**Fig. 7A-B**). However, ZD7288 (50 μM) had no significant effect on the resting membrane potential (RMP) of healthy mouse colon-innervating DRG neurons (**Fig. 7C**). Further analysis of action potential waveform characteristics revealed no differences in the ascending or descending phases of action potentials between neurons treated with ZD7288 and vehicle controls (**Fig. 7D-E**).

**Figure 7.**
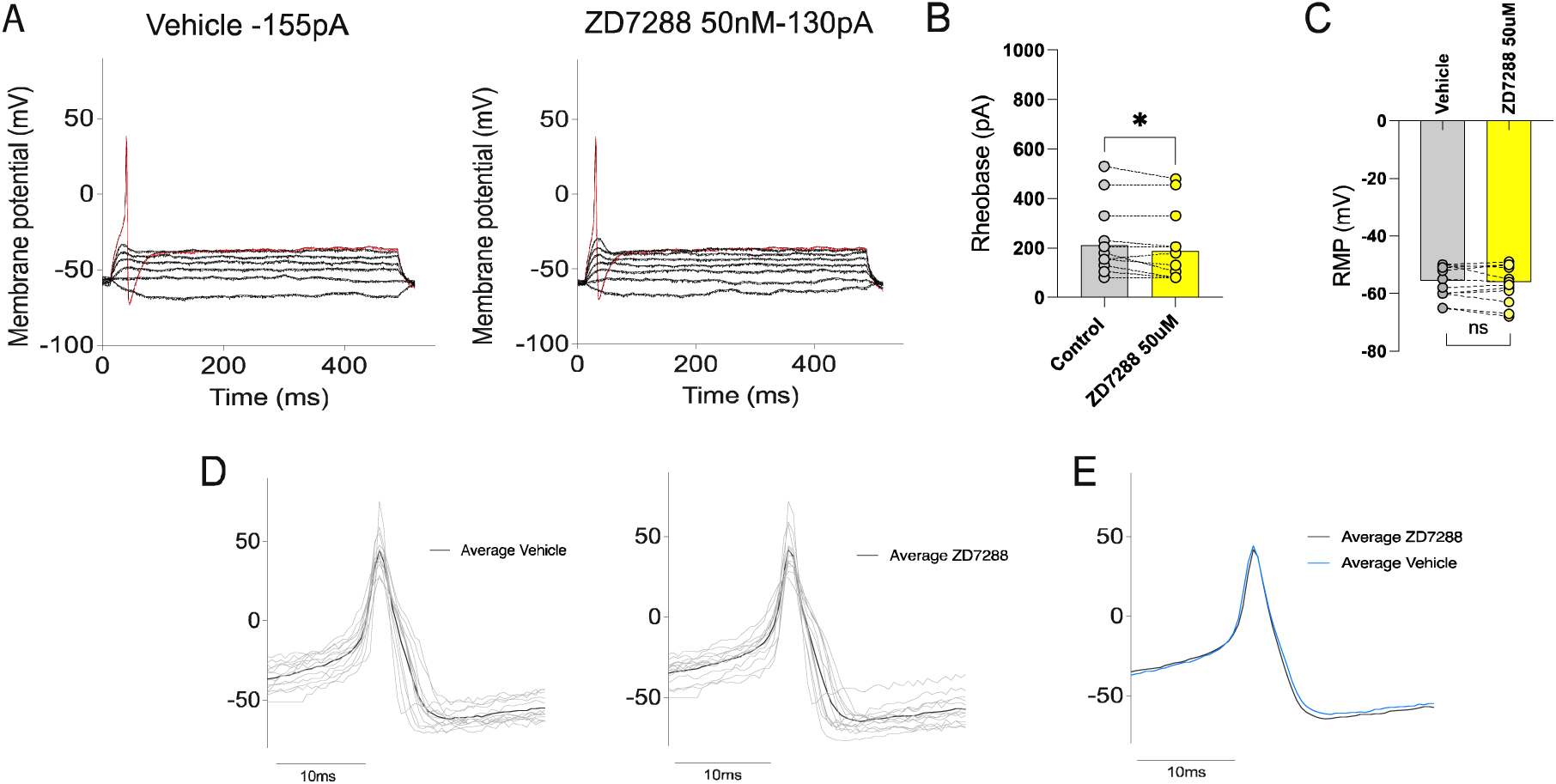
The HCN channel blocker ZD7288 enhances the excitability of mouse colon-innervating DRG neurons. **(A)** Representative voltage responses to current clamp steps recorded from a colon-innervating DRG neuron in the absence (control, vehicle) and presence of ZD7288 (50 μM). Rheobase is indicated in red. **(B, C)** Bar graphs showing the effects of ZD7288 on rheobase and resting membrane potential (RMP) in colon-innervating DRG neurons. Data are presented as mean ± SEM. P values were determined using a paired t-test. **(D)** Representative individual action potential (AP) traces recorded under each condition (grey), with the average waveform overlaid in black and aligned to the peak depolarisation. **(E)** Overlay of average AP waveforms shows no significant differences between vehicle (blue) and ZD7288 (black). n = 14 neurons from N = 3 mice.

## DISCUSSION

GABA_B_Rs are widely expressed throughout the somatosensory nervous system, where they play a key role in modulating pain perception and transmission (43, 44). Targeting GABA_B_Rs has been shown to attenuate mechanical allodynia in neuropathic pain models (45) and reduce colonic nociception, thereby alleviating visceral pain (8). As G-protein-coupled receptors, GABA_B_Rs mediate their inhibitory effects through the modulation of downstream ion channels (46, 47). Notably, these downstream effectors are agonist-specific, with distinct channel interactions depending on the activating ligand (48). For example, the activation of GABA_B_Rs by the agonist baclofen inhibits three distinct calcium channels, Ca_V_2.1, Ca_V_2.2, and Ca_V_2.3, whereas α-conotoxin Vc1.1 activation of GABA_B_Rs selectively inhibits Ca_V_2.2 and Ca_V_2.3 channels, with no effect on Ca_V_2.1 (49). Additionally, GABA_B_R activation by Vc1.1 has been shown to activate G protein-gated inwardly rectifying potassium (GIRK) channels in HEK293T cells co-expressing either heteromeric human GIRK1/2 or homomeric GIRK2 subunits with GABA_B_R (32). This mechanism is particularly relevant given the widespread expression of GIRK channels in neurons throughout the central nervous system, including the spinal cord and sensory ganglia, where they regulate neuronal excitability and contribute to the pathophysiology of several neurological disorders (50, 51). GIRK channels hyperpolarize neurons in response GPCR activation, modulating neuronal activity via GIRK-mediated self-inhibition, slow synaptic potentials, and volume transmission (51).

Notably, activation of GIRK channels via GABA_B_R has been shown to modulate nociceptive transmission (52–55). In mouse DRGs, Vc1.1-induced GABA_B_R activation potentiates GIRK channels, leading to membrane hyperpolarization and decreased neuronal excitability (56). In the present study, we characterized the expression profiles of Ca_V_ and GIRK channels, as well as GABA_B_R subunits, in both mouse and human whole DRG. A recent study utilizing single-cell nucleus RNA sequencing data from human DRGs reported high expression levels of Ca_V_ channels and GABA_B_Rs (57). These findings align with our own, which similarly identified the expression of GABA_B_Rs and Ca_V_s, and demonstrated their functional interactions. We also observed significant expression of GIRK1 channels in human DRGs, with minimal or no expression of other GIRK isoforms, which is consistent with findings from the single-cell nucleus RNA sequencing data of individual human DRG neurons (**Supp. Fig. 1**) (57).

Additionally, we provide evidence that α-conotoxin Vc1.1 inhibits neuronal excitability in mouse DRG through two distinct mechanisms: inhibition of HVA N-type calcium (Ca_V_2.2) channels and potentiation of GIRK1/2 channels. Inhibiting GIRK1/2 channels enhanced neuronal excitability and attenuated the ability of Vc1.1 to suppress action potential firing. Collectively, these findings confirm that in mouse sensory DRG neurons, α-conotoxin Vc1.1 reduces neuronal excitability via GABA_B_R-mediated Ca_V_2.2 channel inhibition and GIRK channel potentiation (**Figure 8**) (32, 56).

**Figure 8.**
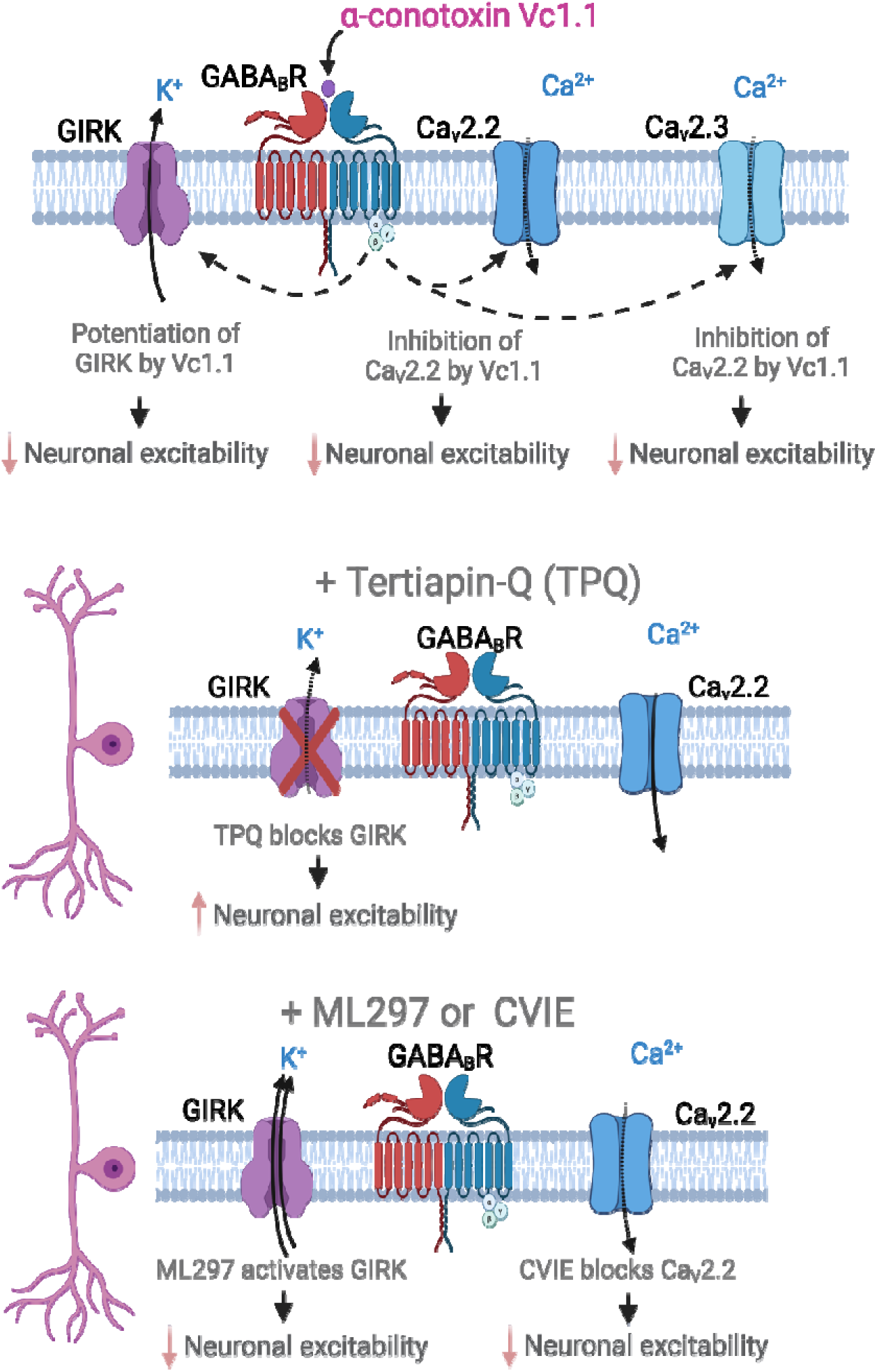
Summary of the mechanisms by which α-conotoxin Vc1.1 inhibits mouse sensory neurons. Whole-cell patch-clamp recordings from mouse dorsal root ganglion (DRG) neurons show that α-conotoxin Vc1.1 significantly reduces neuronal activity by inhibiting Ca_V_2.2 and Ca_V_2.3 channels and potentiating G protein-coupled inwardly rectifying potassium (GIRK) channels, demonstrating its dual role in modulating sensory neuron excitability. Inhibition of GIRK channels with the antagonist Tertiapin-Q (TPQ) increased neuronal excitability, but this effect was not observed in DRG neurons that innervate the mouse colon. In ‘general’ mouse DRG neurons, inhibition of high voltage-activated calcium channels (Ca_V_2.2) by the Ca_V_2.2 antagonist CVIE and activation of GIRK channels by the agonist ML297 both contribute to reduced neuronal excitability. Figure created with Biorender.

Our findings provide important new insights into the downstream coupling of ion channels to GABA_B_Rs. Previous studies have shown that Vc1.1 enhances inwardly-rectifying K^+^ currents in HEK293T cells expressing human GIRK1/2 channels and GABA_B_Rs (32). However, more recent studies in human pluripotent stem cells (hPSC)-derived sensory neurons revealed that while GIRK channels modulate excitability, they do not appear to be functionally coupled to GABA_B_Rs in these cells (31). In the present study, we show GIRK channels are expressed in colon-innervating DRG neurons, with GIRK1 being most prevalent (58% of neurons), followed by GIRK2 (15%), whereas GIRK3 or GIRK4 were not detected. Notably, there was substantial overlap in the expression profile of GABA_B_R, Ca_V_2.2, and GIRK1 channels. However, our afferent nerve recordings and patch-clamp studies showed that GIRK channel inhibition had little impact on colonic afferent function in either healthy or CVH mice. This suggests that, in mouse colonic DRGs, GIRK channels may not play a significant role in regulating neuronal activity.

These findings contrast with previous studies that implicated GIRK channels in modulating neuronal excitability across general DRG populations. This discrepancy aligns with prior research indicating that colon-innervating DRG neurons exhibit distinct functional and transcriptional profiles compared with DRG neurons that innervate other tissues (58–60). One potential explanation is that GABA_B_R-GIRK coupling is inactive or dormant under healthy conditions but becomes functional only during inflammatory or pathological states. Similar context-dependent receptor-effector coupling has been reported for κ-opioid and oxytocin receptors, which mediate anti-nociceptive effects in colonic nociceptors only during inflammatory or post-inflammatory CVH states, but not in healthy conditions (37, 39, 40, 61, 62). However, our patch clamp recordings from colon thoracolumbar DRG neurons in CVH states showed that inhibiting GIRK channels with Tertiapin-Q did not alter rheobase, further supporting the notion that GIRK channels do not significantly regulate excitability in high-threshold nociceptors from the splanchnic pathway or in LT, WDR, or HT pelvic afferents in healthy states.

The analgesic effects of α-conotoxin Vc1.1 on visceral pain are well-established (8, 28–30). Vc1.1-induced activation of GABA_B_R has been shown to inhibit Ca_V_2.2 and Ca_V_2.3 channels, thereby reducing colonic DRG neuronal output by increasing rheobase, decreasing colonic afferent excitability, dampening nociceptive signalling in the spinal dorsal horn in response to colonic distension, and alleviating visceral pain *in vivo* (8, 29, 30). Notably, the anti-nociceptive effects of Vc1.1 are enhanced in hyperalgesic states, as demonstrated in a CVH model, where upregulation of the Ca_V_2.2 exon 37a splice variant has been reported (8). While our data did not support a functional role for GIRK channels in colonic afferents, we identified a significant role for HCN channels in regulating colonic DRG neuronal excitability.

HCN channels are key regulators of neuronal excitability and have been implicated in both somatic (63) and visceral (64) pain. HCN channels are activated by membrane hyperpolarization, conduct both Na^+^ and K^+^ ions, and are modulated by cyclic AMP (cAMP), which facilitates channel opening (65),(66). This is particularly relevant in the context of GABA_B_R signalling, as cAMP-dependent protein kinase phosphorylation has been shown to influence GABA_B_R-effector coupling (67). Using the HCN channel blocker ZD7288, we demonstrated that inhibiting HCN channels altered neuronal excitability in healthy states without significantly affecting membrane potential or action potential properties. This is consistent with transcriptomic studies showing widespread expression of HCN1-4 isoforms in colonic afferent DRG neurons (**Supp. Fig. 2)** (58). Further research is required to determine whether GABA_B_Rs directly interact with HCN channels in colonic DRGs and whether Vc1.1 activation of GABA_B_R modulates HCN channel activity in both healthy and disease states.

In summary, our findings confirm that in sensory mouse DRG neurons, α-conotoxin Vc1.1 reduces neuronal excitability downstream of GABA_B_R activation through both Ca_V_2.2 channel inhibition and GIRK channel potentiation. However, in colonic-innervating DRG neurons, despite strong expression of GIRK1, its functional contribution to afferent responses to mechanical distension or action potential generation was minimal. Instead, HCN channels emerge as promising candidates for modulating excitability in these colon-innervating neurons, warranting further investigation.

## ABBREVIATIONS LIST

CVH: chronic visceral hypersensitivity
DRG: dorsal root ganglion
GABA_B_R: G protein-coupled γ-aminobutyric acid type B receptor
GIRK: G protein-coupled inwardly rectifying potassium
GPCRs: G protein-coupled receptors
HCN: hyperpolarization-activated cyclic nucleotide-gated
hPSC: human pluripotent stem cell
IBS: irritable bowel syndrome
TPQ: tertiapin-Q

## ADDITIONAL INFORMATION SECTION

### Data availability statement

The data that support the findings of this study are available from the corresponding authors upon reasonable request.

### Author contributions

**<u>SAHMRI:</u> M.B**: acquisition, analysis and interpretation of colonic afferent data and patch clamp data from colonic DRG neurons, drafting manuscript and figures, revisions for important intellectual content. **S.G.C**: acquisition, analysis and interpretation of QPCR from mouse and human DRG and single-cell RT-PCR from colonic DRG neurons, drafting manuscript and figures, revision for important intellectual content. **S.M.B**: conceptualization of colonic studies, interpretation of data, project administration, funding acquisition, writing, review and editing, resources, methodology, supervision. **<u>University of Wollongong:</u> A.R.B**: acquisition, analysis and interpretation of patch clamp data from adult mouse DRG neurons, drafting manuscript and figures, revisions for important intellectual content. **D.J.A**: conceptualization, interpretation of data, project administration, funding acquisition, writing, review and editing, resources, methodology, supervision.

All authors approved the final version of the manuscript; agree to be accountable for all aspects of the work in ensuring that questions related to the accuracy or integrity of any part of the work are appropriately investigated and resolved; and all persons designated as authors qualify for authorship, and all those who qualify for authorship are listed.

