## Supplementary material for "Differential contributions of Ca_V_2.2, GIRK, and HCN channel to the modulation of excitability by α-conotoxin Vc1.1 and baclofen in somatic and visceral sensory neurons": Supp Info

**Running Title:** Roles of Ca_V_2.2 and GIRK channels in DRG neurons

Mariana Brizuela ^1^, Anuja R. Bony ^3^, Sonia Garcia Caraballo ^1^, David J. Adams ^3,*^, and Stuart M. Brierley ^1,2,*^

^1^ Visceral Pain Research Group, South Australian Health and Medical Research Institute (SAHMRI), North Terrace, Adelaide, SA 5000 Australia.

^2^ Faculty of Health and Medical Sciences, University of Adelaide, North Terrace, Adelaide, South Australia 5000, Australia.

^3^ Molecular Horizons/Faculty of Science, Medicine and Health, University of Wollongong, Wollongong, NSW 2522 Australia.

***Corresponding authors:**

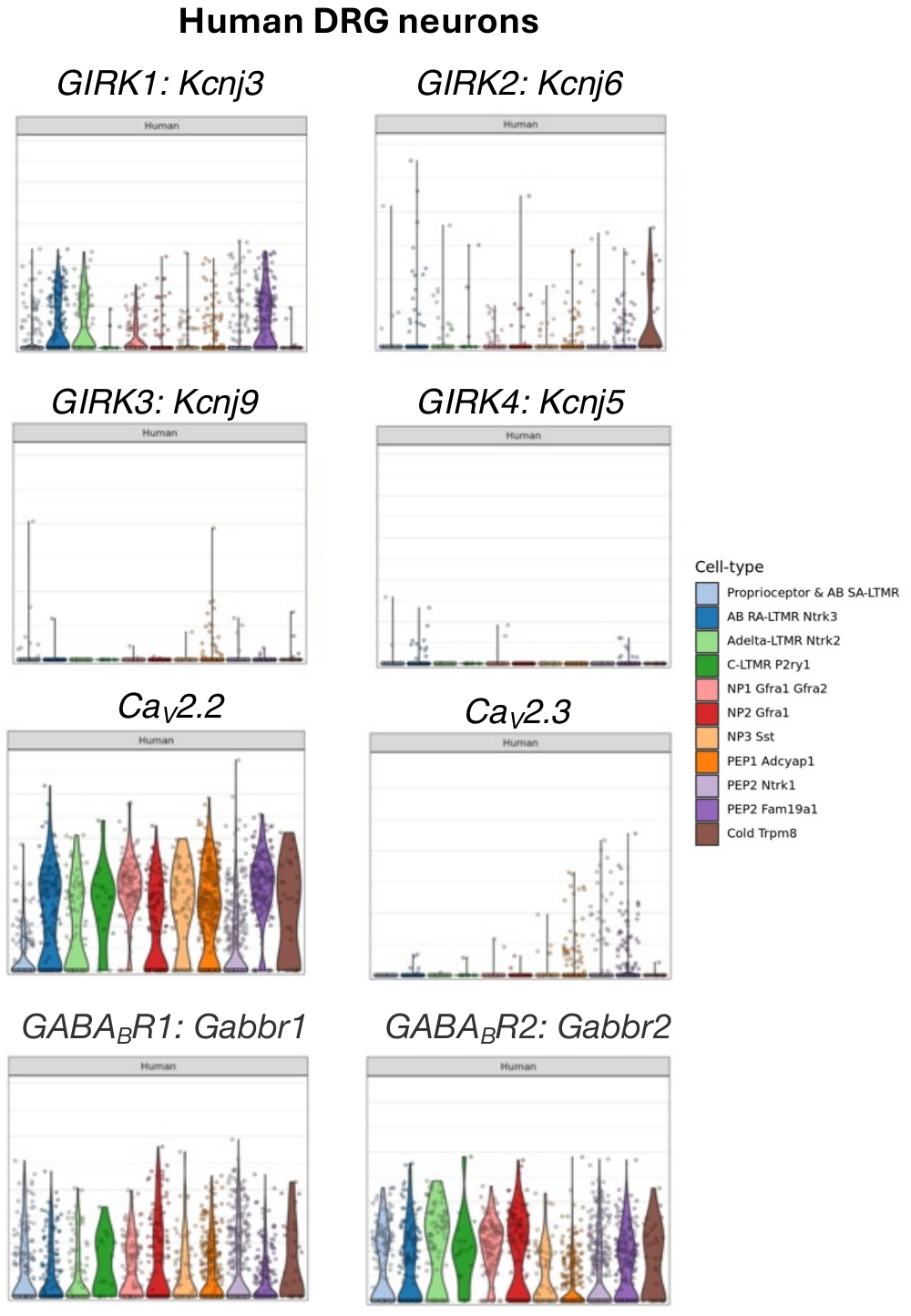

**Supplementary Figure 1: Expression of GIRK, Ca_V_2.2, Ca_V_2.3, and GABA_B_R1 and GABA_B_R2 in individual human DRG neurons based on single-cell transcriptomics.**

Graphs were generated for comparative purposes by using the website <http://research-pub.gene.com/XSpecies>, which presents data from the study by Jung *et al.,* *“Cross-species transcriptomic atlas of dorsal root ganglia reveals species-specific programs for sensory function”* (1). Gene-specific expression profiles were obtained by inputting the relevant genes of interest into the platform.

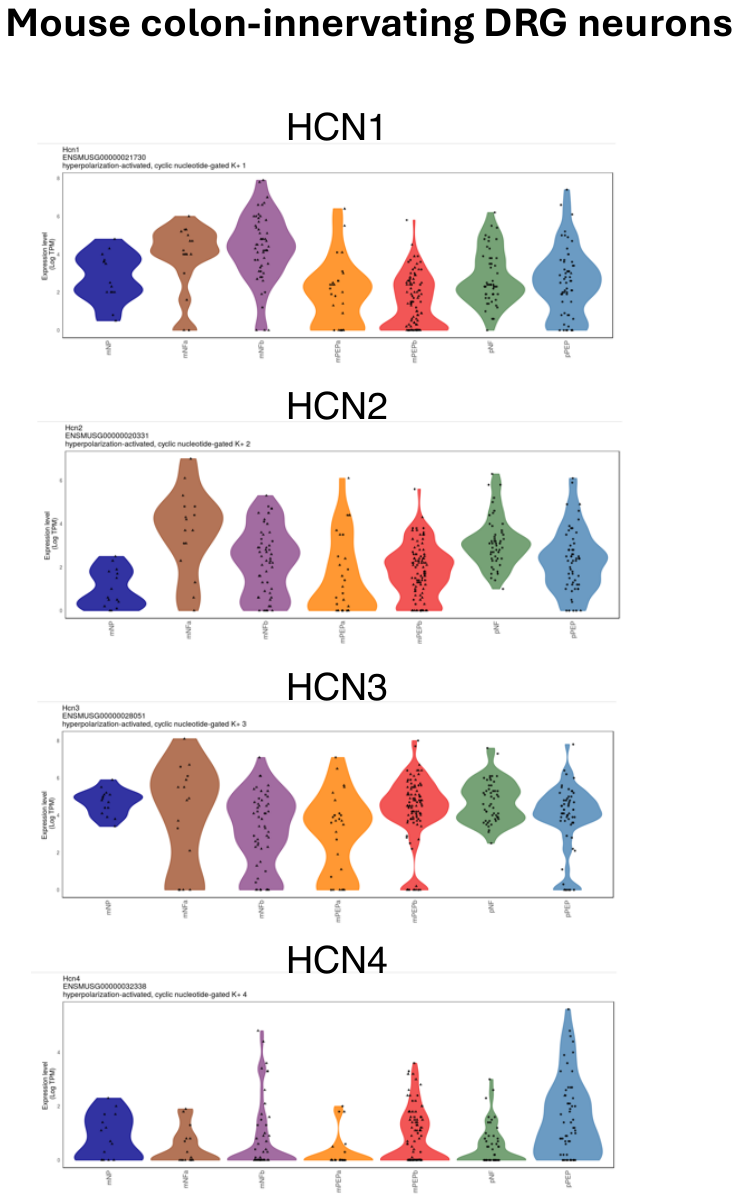

**Supplementary Figure 2: Expression of HCN1-4 in individual DRG neurons innervating the mouse colon based on single-cell transcriptomics data.**

Graphs were generated for comparative purposes using the website <http://hockley.shinyapps.io/ColonicRNAseq>, which presents data from the study by Hockley *et al.,* *“Single-Cell RNAseq Reveals Seven Classes of Colonic Sensory Neuron”* (2). Gene-specific expression profiles were obtained by inputting the respective genes of interest into the platform.

**Supplementary Information References**

1.	**Jung M, Dourado M, Maksymetz J, Jacobson A, Laufer BI, Baca M, Foreman O, Hackos DH, Riol-Blanco L, and Kaminker JS**. Cross-species transcriptomic atlas of dorsal root ganglia reveals species-specific programs for sensory function. *Nature Communications* 14(1): 366, 2023. doi: 10.1038/s41467-023-36014-0.

2.	**Hockley JRF, Taylor TS, Callejo G, Wilbrey AL, Gutteridge A, Bach K, Winchester WJ, Bulmer DC, McMurray G, and Smith ESJ**. Single-cell RNAseq reveals seven classes of colonic sensory neuron. *Gut* 68(4): 633–644, 2019. doi: 10.1136/gutjnl-2017-315631.
